# Hepatitis E virus genome replication is independent of cyclophilins A and B

**DOI:** 10.1101/2022.12.05.519105

**Authors:** Frazer J.T. Buchanan, Shucheng Chen, Mark Harris, Morgan R. Herod

**Affiliations:** School of Molecular and Cellular Biology, Faculty of Biological Sciences and Astbury Centre for Structural Molecular Biology, University of Leeds, Leeds, LS2 9JT, UK

**Keywords:** HEV, replication complex, replicon, zoonosis, cyclosporine

## Abstract

Hepatitis E virus (HEV) is an emerging pathogen responsible for more than 20 million cases of acute hepatitis globally per annum. Healthy individuals typically have a self-limiting infection, however, mortality rates in some populations such as pregnant women can reach 30%. A detailed understanding of the virus lifecycle is lacking, mainly due to limitations in experimental systems. In this regard, the cyclophilins are an important family of proteins that have peptidyl-prolyl isomerase activity and play roles in the replication of a number of positive-sense RNA viruses, including hepatotropic viruses such as hepatitis C virus (HCV). Cyclophilin A (CypA) and cyclophilin B (CypB) are the two most abundant human cyclophilins in hepatocytes and are therefore potential targets for pan-viral therapeutics. Here, we investigated the importance of CypA and CypB for HEV genome replication using a sub-genomic replicon system. This system removes the requirements for viral entry and packaging and therefore allows for the sensitive measurement of viral genome replication in isolation. Using pharmacological inhibition by cyclosporine A (CsA), known to suppress HCV replication, and silencing by shRNA we find that CypA and CypB are not essential for replication of genotype 1 or 3 HEV replication. However, we find that silencing of CypB reduces replication of genotype 1 HEV in some cells, but not genotype 3. These data suggests HEV is atypical in its requirements for cyclophilin for viral genome replication and that this phenomenon could be genotype specific.

## Introduction

Hepatitis E virus (HEV) is one of the leading etiological agents of acute hepatitis and is responsible for more than 20 million cases annually. The virus is a member of the *Orthohepevirus* genus within the *Hepeviridae* family. The genus is divided into 4 species groups (A-D), which can infect a wide range of animals, including humans, and are classified into 7 genotypes (G1-G7) [1, 2]. Genotype 1 and 2 viruses appear to be obligate human pathogens that are transmitted between humans by the faecal-oral route, with the potential to cause large outbreaks [2, 3]. Genotype 3 and 4 viruses have been isolated in several animal species including humans and are of particular concern as they have been associated primarily as a porcine zoonosis in higher and middle-income countries [4, 5]. Infection in healthy individuals usually leads to acute hepatitis which has a low rate of mortality. However, infection during pregnancy is of particular concern as mortality rates have been reported to be up to 30% [6]. This higher risk of mortality has also been observed in immunocompromised individuals. No specific regimen of treatment is used to treat infected individuals, with antivirals such as ribavirin used in combination with supportive care [7].

HEV is a single-stranded positive-sense RNA virus with a genome length of approximately 7.2 kb. The genome contains three open reading frames (ORFs). ORF1 is translated to produce the viral polyprotein that contains the protein domains required for viral RNA replication. The second and third open reading frames, ORF2 and ORF3, are translated into the viral capsid protein and a small membrane protein involved in virus release, respectively [7, 8]. A fourth open reading frame, ORF4, has also been identified but only in genotype 1 viruses [9]. Replication of the viral genome is controlled by the ORF1 polyprotein, also known as pORF1. Through sequence homology to related virus families such as the caliciviruses and togaviruses, pORF1 has been predicted to contain at least six distinct protein domains. At the N-terminus of the polyprotein is a methyltransferase (MeT) domain, which is followed by a putative cysteine protease (PCP) domain. Spanning the centre of the polyprotein is a stretch of high sequence diversity, termed the hyper variable region (HVR), and followed by the X domain that is hypothesised to contain a macro domain. At the C-terminus of the polyprotein are the putative viral helicase (Hel) and RNA-dependant RNA-polymerase (RdRp). Based on considerable sequence homology the function of the MeT, Hel and RdRp, domains are highly probable and the MeT, Hel and X domains have been formally shown to have a biochemical function [10, 11, 12, 13, 14]. However, some regions, such as the HVR have poor sequence homology and no function has been suggested.

HEV is a hepatotropic virus with hepatocytes being the primary site of infection and pathology. Small molecules that inhibit the replication of hepatotropic viruses such as hepatitis C virus (HCV), have shown promise as therapies as well as tools for understanding fundamental virus biology. The cyclophilins (Cyps) are a family of peptidyl prolyl isomerases that aid in a number of cellular processes such as protein folding, trafficking and innate immune signalling, and have been identified as proteins that are co-opted by viruses to promote their replication. Cyclophilin A (CypA) is the predominant human cyclophilin which has been documented to be important for the replication of a number of viruses including SARS coronavirus, HIV and HCV [15, 16]. In the case of HCV, CypA is bound by the viral non-structural protein NS5A, to directly inhibit protein kinase R (PKR) and prevent interferon expression. HCV is thought to use this mechanism to help in evasion of the innate immune system (Daijun et al., 2012, Fernandes et al., 2010). Furthermore, inhibition of CypA with the selective CypA inhibitor cyclosporine A (CsA) has been used medically to prevent organ transplant rejection [17]. Additionally, it has been reported that cyclophilin B (CypB) is important for HCV replication [15].

The literature regarding the role of the Cyps in HEV infection is currently divided. Wu et al [18] reported that CypA disruption did not impact the replication of HEV primary isolates in cell culture. Contrary to this, Wang et al [19] reported that native functional CypA inhibits replication of HEV in hepatocytes, and that inhibition of CypA with CsA actually promotes HEV replication. Given the importance of CypA and CypB in the replication of other chronic hepatotropic viruses, they represent possible pan-therapeutic targets. We therefore decided to investigate thoroughly a potential role for these proteins in HEV replication. Using sub-genomic replicons (SGR) of HEV, we compared the effect of CypA and CypB pharmacological and genetic inactivation on viral genome replication. Using this approach our data suggests that CypA or CypB or not essential for genotype 1 or genotype 3 HEV replication. However, silencing of CypB can reduce replication of genotype 1 HEV in some cell types. These data suggest that in some cells CypB may have an auxiliary role in HEV replication.

## Material and Methods

### Cell lines and plasmids

Huh7, Huh7.5 and HEK293T cells were maintained in Dulbecco’s modified Eagle’s medium with glutamine (Sigma-Aldrich) supplemented with 10 % FCS, 1 x non-essential amino acids (Gibco) 50 U / mL penicillin and 50 μg / mL streptomycin (Sigma-Aldrich).

A plasmid carrying the wild-type genotype 1 HEV replicon expressing GFP, pSK-E2-GFP, was a kind gift from Dr Patrizia Farci and has been described previously [20]. This plasmid was modified to replace the GFP open reading frame with nano-luciferase as previously described [21] to produce pSK-E2-nLuc. Mutations within these plasmids were performed by standard two-step overlapping PCR mutagenesis. Negative control replicons were generated containing a double point mutation in the RdRp active site GDD motif (GNN) and has been previously described [20] Plasmid carrying wild-type genotype 3 HEV replicon expression nano-luciferase, pUC-HEV83-2, was a kind gift from Dr Koji Ishii and has been described previously [22, 23].

### Generation of silenced cell lines

HEK293T cells were prepared for transfection in 10 cm dishes. Lentivirus production was initiated via transfection of the following into HEK293T cells; 1 μg p8.9 (packaging plasmid) 1 μg pMDG (VSVg envelope plasmid) 1.5 μg pHIV-SIREN encoding shRNA (genome plasmid). Supernatants were harvested at 48 h and 72 h post-transfection. Supernatants were filtered (0.45 μm) and stored at -80°C.

Huh7 or Huh7.5 cells were plated at a density of 1 × 10^5^cells / well in a 6-well plate. Cells were then transduced with 1 mL / well of lentivirus supernatant in the presence of 8 μg / mL polybrene to promote transduction. Selection with 2.5 μg / mL puromycin was introduced 72 h post-transduction. Passage of cells in puromycin selection media was continued to maintain expression of shRNA.

### *In vitro* transcription

pSK-E2-nLuc replicon plasmid was linearised with *Bgl*II and pUC-HEV83-2 replicon plasmid was linearised with *Hind*III before being used to generate T7 *in vitro* transcribed RNA using the HiScribe T7 ARCA mRNA kit with tailing following manufacturer’s instructions (Promega). RNA was purified using an RNA clean and concentrate kit (Zymo Research) and the quality was checked using a MOPS/formaldehyde agarose gel electrophoresis.

### Replication assays

Replicon experiments were conducted as previously described (Herod et al., 2022). Briefly, Huh7 or Huh7.5 cells were detached by trypsin, washed twice in ice-cold DEPC-treated PBS and re-suspended at 1 × 10^7^ cells / mL in DEPC-treated PBS. Subsequently 400 μL of cells was mixed with 2 μg of RNA transcript, transferred to a 4 mm gap electroporation cuvette (SLS) and pulsed at 260 V, 25 ms pulse length in a Bio-Rad Gene Pulser (Bio-Rad) on the square wave setting. Electroporated cells were recovered into 4 mL media, seeded into replicate 6-well tissue culture plates, and replication measured at 24 h intervals using the Nano-Glo luciferase assay system (Promega). For cyclosporine A treatment the electroporated cells were seeded into replicates of 24-well plates, allowed to adhere before the media was replaced with fresh media containing cyclosporine (all Sigma-Aldrich), at the indicated concentration.

### MTS assay

The cell viability experiments were conducted by seeding cells into 96-well plates, allowing to adhere for 24 h before addition of a serial dilution of inhibitor and measurement of cell viability 72 h later using the CellTiter AQueous One solution (Promega), following manufacturer’s instructions. Briefly, 20 μL of reagent was added to each well before samples and appropriate media only blanks were incubated for 45-60 mins at 37°C. Absorbance at 490 nm was measured on an Infinite F50 (Tecan).

### Western blotting

Cell lysates were centrifuged for 20 mins at 17,000g, supernatant removed to a separate tubes and mixed with an equal volume of 2x Laemmli buffer (Sigma-Aldrich). Samples were heated for 5 mins at 100°C and separated on a 10 % sodium dodecyl sulphate polyacrylamide gel. Proteins were transferred onto Immobilon transfer membrane (Merck) using a BioRad Trans-Blot turbo transfer system. Membranes were blocked in 10 % milk in tris-buffered saline solution containing 0.1 % Tween (Fisher). Membranes were then incubated overnight at 4°C with rabbit anti-CypA (1:1000) (Enzo) or anti-CypB (1:2000) (Abcam) antibody. Membranes were washed three times prior to 1 h incubation with anti-rabbit horse radish peroxidase conjugated secondary antibody. Membranes were washed three times and incubated in ECL reagent (Thermo scientific) before exposure to CL-Xposure film (Thermo scientific), and developed by XOgraph (Fuji).

## Results

### Pharmacological inhibition of cyclophilin does not impact HEV genome replication

Previous work has established that functional CypA is necessary to support the replication of several positive-sense RNA viruses, such as HCV [15, 16, 24]. However, the role for cyclophilins in the replication of HEV remains disputed, in part due to the difficulty in investigating separate parts of the viral replication cycle in isolation. To elucidate the effects of Cyps on HEV genome replication we employed an HEV sub-genomic replicon (SGR) a self-replicating RNA in which a portion of the viral structural proteins are replaced by a nano-luciferase (nLuc) reporter gene (Figure 1). Measurement of nLuc activity allows for an indirect measure of viral genome replication in the absence of virus entry or assembly. Cyclosporine A (CsA) is a potent inhibitor of both cyclophilin A and cyclophilin B. It is a cyclic molecule derived from the fungus *Tolypocladium inflatum* and complexes with cyclophilin to prevent them carrying out catalytic peptidyl prolyl isomerisation as well as preventing interactions with other cellular proteins [25, 26]. We therefore decided to start by investigating the sensitivity of HEV replication to CsA.

**Figure 1.**
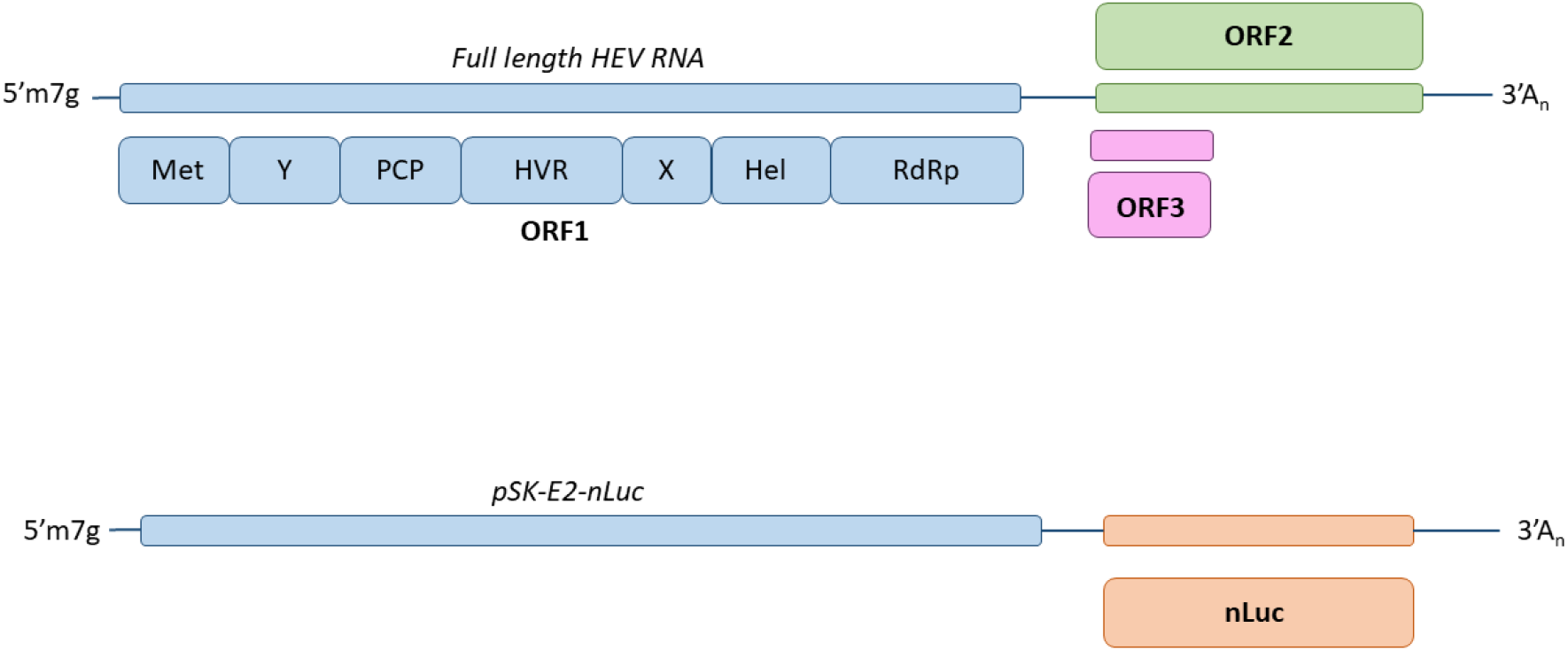
HEV genome organisation vs replicon. Schematic of the HEV genome showing open reading frames 1-3 (ORF1-3). ORF1 is reported to contain a methyltransferase (Met), Y domain (Y), putative cysteine protease (PCP), hyper variable region (HVR), macro domain (X), helicase domain (Hel) and RNA dependent RNA polymerase (RdRp). ORF2 and ORF3 are both produced from a viral subgenomic RNA. Nano-luciferase replicon pSK-E2-nLuc was created by replacing ORF2 and ORF3 with nano-luciferase (nLuc) to act as a reporter for replication.

Two human hepatocellular carcinoma lines that support HEV replication (Huh7 and the derivative cell line Huh7.5, which contain a RIG-I mutation and support improved replication of viruses such as HCV [27, 28]) were transfected with a genotype 1 HEV SGR RNA (SKE2nLuc), which is derived from the Sar55 infectious clone sequence [20] (Figure 1). Alongside this, cells were also transfected with an equivalent replication defective SGR (SKE2nLuc-GNN), which contained two inactivating mutations in the active site of the viral RNA polymerase. CsA was added to the growth medium 24 h after transfection at varying concentrations (0 - 100 μM), and replication assayed daily for 120 h post-transfection (Figure 2).

**Figure 2.**
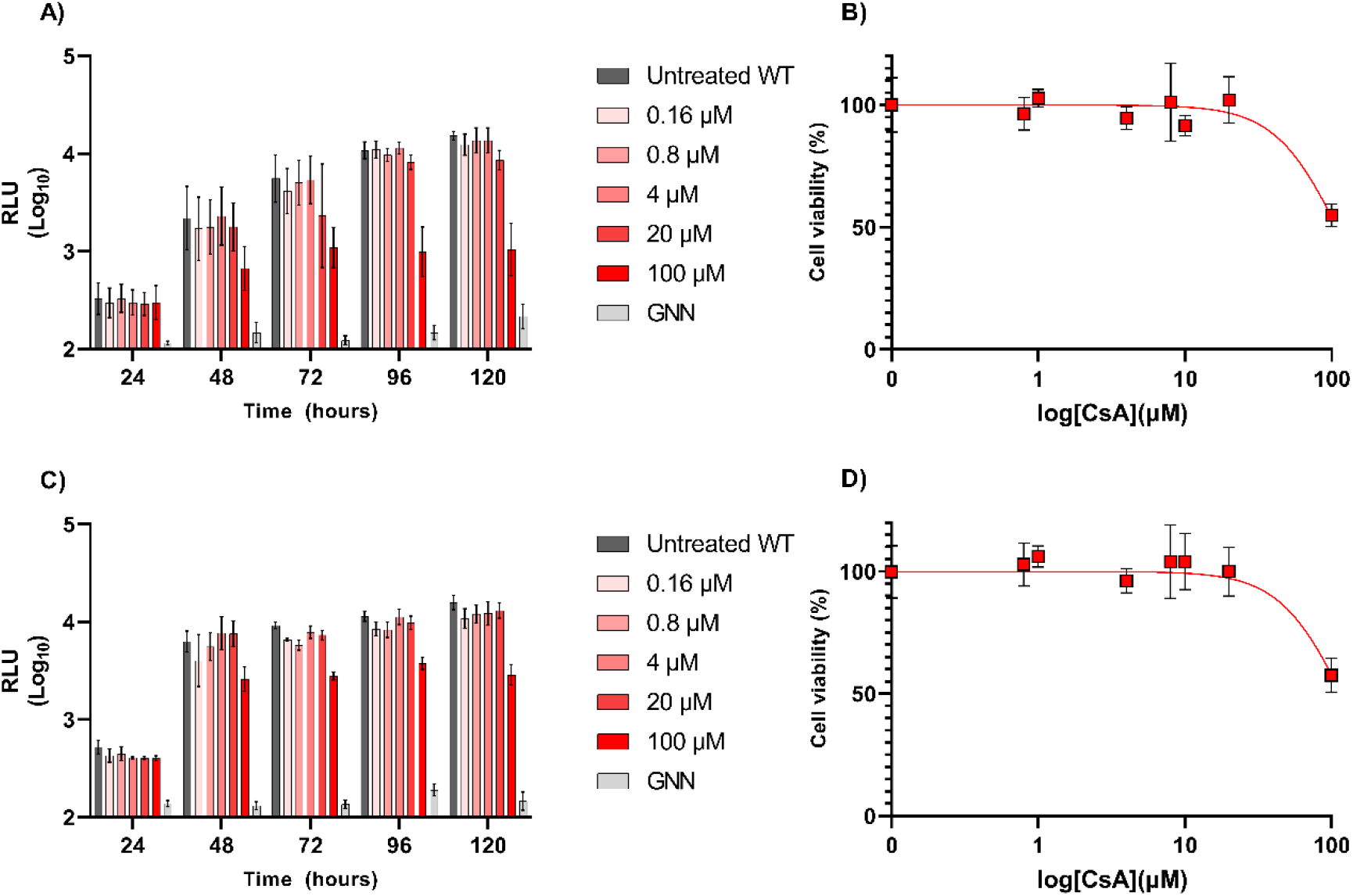
CsA dose response in HEV transfected hepatocytes. **A)** Huh7 cells or **C)** Huh7.5 cells were electroporated with wild-type (WT) SKE2nLuc SGR or GNN SGR RNA prior to addition of CsA at varying concentrations (0-100 μM) 24 h post-electroporation. Cells were harvested at 24 h intervals for 120 h and luciferase activity determined. Data are presented as mean luciferase activity as relative light units (RLU) (n= 3 +/-SEM). **B)** Huh7 or **C)** Huh7.5 were seeded into 96-well plates, allowed to adhere for 24 h before replicate wells were treated with a serial dilution of cyclosporine (0 – 100 μM). Replicate wells were left untreated or treated with DMSO solvent only as controls. 72 h after treatment cell viability of Huh7 cells was calculated by MTS assay. Data presented as mean percentage cell viability, normalised to untreated controls (n = 3 +/-SEM).

For the wild-type (WT) untreated SGR, nLuc activity increased approximately 100-fold over the duration of the experiment. As anticipated, the replication defective replicon (GNN) only demonstrated background levels of nLuc activity at every time point. In comparison to the untreated WT SGR there was no marked difference in nLuc activity upon treatment of CsA up to a concentration of 20 μM. In contrast, there was approximately a 11-fold decrease in nLuc activity in cells treated with 100 μM of CsA compared to untreated cells by day 5 post-electroporation. The pattern of results remained consistent in Huh7 cells when compared to Huh7.5 cells, suggesting both cell lines are able to support replication to a similar level. These data suggest only the highest concentration of CsA used (100 μM) reduced luciferase activity. However, concentrations of CsA above 20 μM are reported to be cytotoxic [29, 30]. To quantify any difference between cytotoxicity and inhibition of replication we conducted comparative cytotoxicity experiments. Huh7 and Huh7.5 cells were treated with a serial dilution of CsA and cytotoxicity evaluated by MTS assay three days post-treatment (Figure 2). Cytotoxicity was similar in both Huh7 and Huh7.5 cells with >75% viability at concentrations of 20 μM and under. However, at 100 μM Huh7 and Huh7.5 cells showed average cell viability of ∼56 %, which is similar to the reduction in nLuc activity observed (Figure 2). Taken together these data would suggest that CsA treatment does not reduce HEV replication at sub-cytotoxic concentrations. We conclude from these data that pharmacological inhibition of cyclophilin does not affect HEV replication.

### CypA is not essential for replication of HEV in Huh7 nor Huh7.5 cells

Pharmacological inhibition of Cyps by CsA only suppressed HEV genome replication at cytotoxic concentrations. In order to distinguish isomerase activity from other cellular functions that could be involved in HEV replication, we adopted a genetic approach to silence CypA expression by lentiviral delivery of shRNA in both Huh7 and Huh7.5 cells. We first confirmed silencing of CypA expression by western blot, alongside scramble shRNA controls (Figure 3A). Both Huh7 and Huh7.5 shCypA silenced cell lines produced less CypA compared to the scramble control.

**Figure 3.**
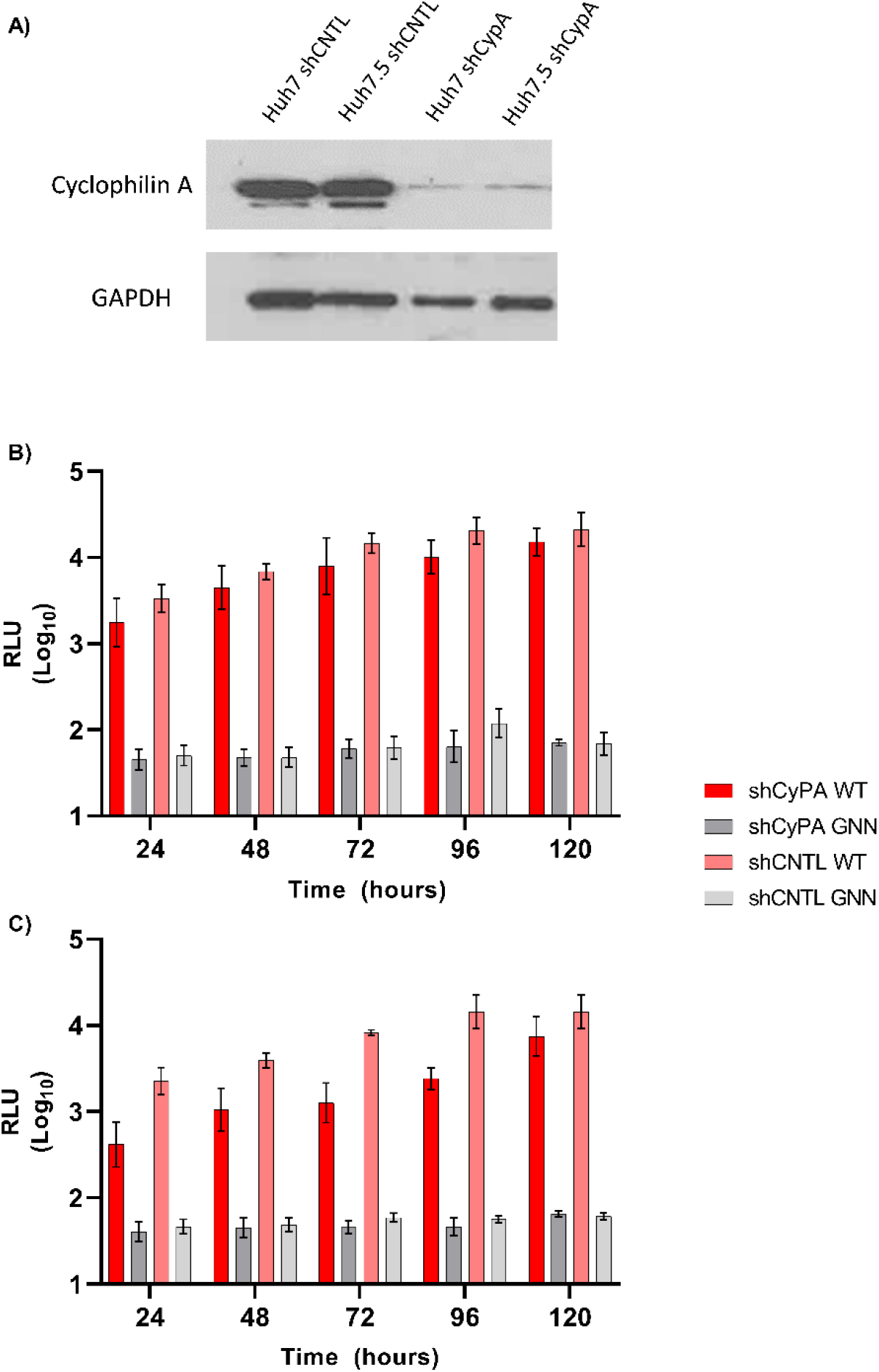
CypA is not essential for HEV replication in Huh7 or Huh7.5 cells. **A)** Detection of CypA expression by western blot in Huh7 and Huh7.5 silenced cell lines (shCypA) and scramble controls (shRNA) with GAPDH used as a loading control. Stable clones of **B)** Huh7 or **C)** Huh7.5 cells silenced for CypA by shRNA (shCypA) or a scramble shRNA control (shCNTL), were electroporated with the wild-type (WT) SKE2nLuc or non-replicating SKE2nLuc-GNN control (GNN) RNA. Cells were harvested at 24 h intervals for 120 h and luciferase activity determined. Data are presented as mean luciferase activity as relative light units (RLU) (n = 3 +/-SEM).

Following validation of reduced CypA expression, the CypA silenced cell lines and scrambled controls were transfected with the SKE2nLuc (WT) replicon RNA or SKE2nLuc-GNN (GNN) control and nLuc activity measured over 120 h post-transfection (Figure 3B & C). Ablation of CypA in Huh7 cells did not reduce replication with nLuc expression equivalent to the scrambled control cell line at every time point of the experiments. There was a minor reduction in replication by five days post-electroporation which was not significant. Silencing of CypA in Huh7.5 cells led to a ∼1.5-fold decrease in nLuc activity 120 h post-electroporation, however this was not significant. There was no significant difference in nLuc expression between the CypA silenced and scramble control cell lines at any other time points. For both experiments, the GNN replicon only produced background levels of luciferase at all time points in both cell types.

### CypB silencing limits HEV replication in Huh7 and Huh7.5 cells

CypA is not the only cyclophilin that is known to be important for viral replication in host cells. CypB has been found to be important for replication in other RNA viruses such as HCV and Japanese Encephalitis virus [31, 32]. After establishing CypA had no essential role in HEV RNA replication we turned our attention to CypB. In order to investigate a role for this protein in HEV replication, we adopted the silencing technique described above. shRNA was used to stably silence expression of CypB in Huh7 and Huh7.5 cells. As before we verified silencing of cyclophilin B by western blot (Figure 4A). Scrambled shRNA sequence was also maintained as a control for both cell types. The CypB silenced cell line and the previously described scrambled control were transfected with the SKE2nLuc (WT) replicon or SKE2nLuc-GNN (GNN) control and nLuc activity measured over 120 h post-transfection as before.

**Figure 4.**
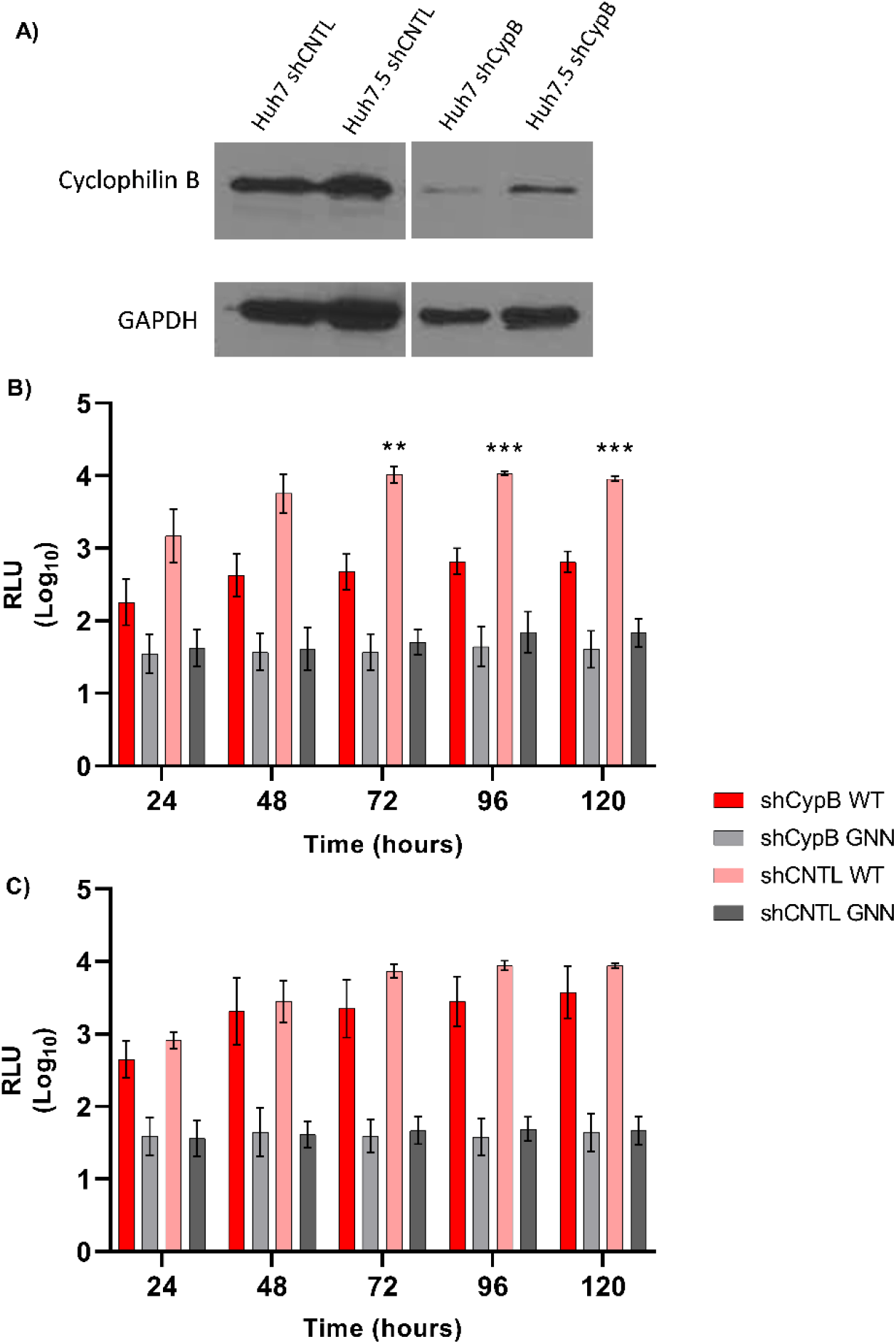
CypB is necessary for efficient HEV replication in Huh7 but not Huh7.5 cells. **A)** Detection of Cyclophilin B expression via western blot in Huh7 and Huh7.5 silenced cell lines (shCypA) and scramble controls (shRNA) with GAPDH used as a loading control. Stable clones of **B)** Huh7 or **C)** Huh7.5 cells silenced for CypB by shRNA (shCypB) or scramble shRNA control (shCNTL), were electroporated with the wild-type (WT) SKE2nLuc RNA or non-replicating SKE2nLuc-GNN control (GNN). Cells were harvested at 24 h intervals for 120 h and luciferase activity determined. Data are presented as mean luciferase activity as relative light units (RLU) (n=3 +/-SEM, *** p-value <0.001; * p-value <0.05).

In contrast to CypA silencing (Figure 2), CypB silencing in Huh7 cells significantly reduced HEV replication between 72 h to 120 h post-electroporation by approximately ∼12-fold compared to the scramble control. There was no significant different in HEV replication in Huh7.5 cells ablated for CypB at any time points compared to the scrambled control. As before, the GNN replicon only produced background levels of luciferase at all-time points in all cell types.

### Genotype specific differences in the requirement for CypA and CypB

The effect of the Cyp proteins on HEV replication has yielded conflicting results, which could potentially be the consequence of variances in the viral genotypes investigated. We noted that Wu et al [18] found that treatment with CsA led to a decrease in total HEV RNA for genotypes 1 and 3 in their cell culture system. In contrast, Wang et al [19] found that ablation of CypA or CypB increased total HEV RNA. It was therefore important to extend our investigations to include other HEV genotypes of human importance such as genotype 3 viruses. Using the CypA or CypB silenced cells (as described in Figures 3 and 4), we investigated the replication of an nLuc containing G3 replicon (HEV-83-2-nLuc) derived from the G3-HEV-83-2-27 infectious clone sequence. All four cell lines together with the shRNA scrambled controls were electroporated with the HEV-83-2-nLuc WT SGR and nLuc activity measured over 120 h post-transfection.

CypA or CypB silencing in Huh7 cells reduced G3 replication by ∼3-fold at 24 h to 120 h post-electroporation, however this was reduction was not statistically significant compared to the scramble control. There was no difference in replication in in Huh7.5 cells ablated for CypA (Figure 5). Likewise, CypB ablation in Huh7.5 cells led to small 2-to 4-fold reduction in luciferase expression at days 72 h to 120 h post-electroporation but this was not statistically significant. We conclude that CypA is not essential for efficient replication of G3 HEV in these cells.

**Figure 5.**
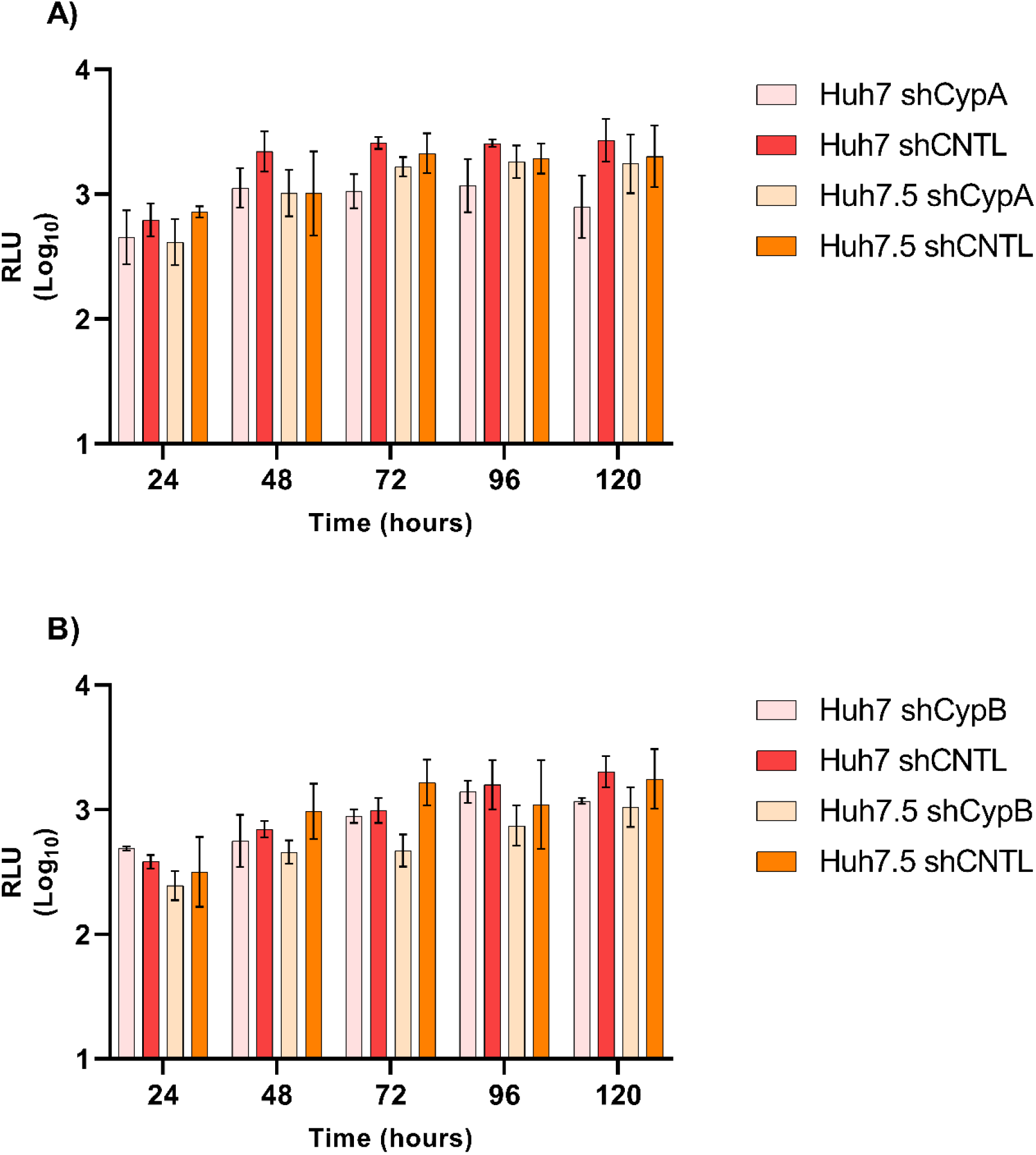
CypA is necessary for efficient HEV Gt-3 replication whilst CypB is not. Stable clones of **A)** Huh7 and Huh7.5 cells silenced for CypA by shRNA (shCypA), or scramble control (shCNTL), were electroporated with the WT genotype 3 SGR HEV-83-2-nLuc. **B)** Huh7 and Huh7.5 cells were silenced for CypB (shCypB), or scrambled control (shCNTL), were electroporated with the WT genotype 3 SGR HEV-83-2-nLuc. Cells were harvested at 24 h intervals for 120 h and luciferase activity determined. Data are presented as mean luciferase activity as relative light units (RLU) (n=3 +/-SEM).

## Discussion

The role of host cell factors in viral propagation is an important aspect of infection. The cyclophilin have been identified as such factors, they can be co-opted by viruses to aid in the completion of their replication cycles and formation of viral particles [15, 16].

HCV, a well-studied hepatotropic virus, relies on the CypA complex in order to evade PKR mediated innate immune signalling during hepatocyte infection, favouring the formation of membrane bound replication sites (the membranous web) via NS5A-mediated inhibition of CypA. Additionally, CypB also complexes with the NS5B polymerase and contributes to genome replication [33]. The tissue tropism of HEV is very similar to that of HCV, hepatocytes being the primary replication site. Despite this, the importance of cyclophilin in HEV replication remains disputed [18, 19].

Wu et al, [18] adopted the use of induced pluripotent stem cell-hepatocyte like cells (iPSC-HLCs) to investigate cell culture and non-cell culture adapted strains of HEV. iPSC-HLCs infected with non-cell culture adapted HEV genotypes 1-4 were treated with CsA. Total RNA was measured as an indication of replication, and CsA did not have an effect on genotype 1. These results agree with our findings, that CsA does not enhance nor impede genotype 1 HEV replication. Interestingly they also found that p6 (a cell culture adapted isolate of genotype 3) showed enhanced HEV replication under CsA treatment. The differences between adapted and non-cell culture adapted strains of HEV informed our experimental design. Thus, we chose two standard human hepatocellular carcinoma cell lines widely used in both HCV and HEV research and two different HEV genotypes to bring consistency into the investigation of the role of cyclophilin in HEV replication.

In contrast to both our study and Wu et al [18], Wang et al [19] found that CsA enhanced replication of the HEV p6 isolate in Huh7 cells in a dose-dependent fashion. Our results also contrast with those of Wang et al [19] as we found that ablation of CypA did not impact genotype 1 HEV replication in hepatocytes. Additionally, we found that CypB ablation led to a modest reduction in genotype 1 HEV replication which disputes the findings of Wang et al that CypB ablation enhances HEV replication. To further clarify the role of the cyclophilin in HEV replication, we repeated our luciferase assays for genotype 3 HEV in CypA and CypB ablated cells. These data demonstrated that neither CypA nor CypB impacted on genotype 3 replication.

The contrasting data reported in this study compared to the studies by Wu et al [18] and Wang et al [19], could be attributed to several differences. Firstly, we considered that differences in genotypes of virus used might explain the discrepancies. Wang et al used a genotype 3 based SGR from a cell culture adapted HEV strain and Wu et al [18] used primary HEV isolated from genotype 1. There is a possibility therefore that the dependence on CypA and or CypB is genotype specific, which requires further investigation using SGRs and infectious virus system in physiologically relevant contexts.

Several RNA viruses require functional CypA in order to complete their replication cycle. However, this requirement is not universal as CypA is dispensable for Chikungunya RNA viral replication [34]. Additionally, replication of hepatitis A virus (HAV) in Huh7 cells has been reported to be independent of CypA [35]. These observations were validated pharmacologically and genetically, like our data here. Potentially similar to these viruses we speculate that HEV is able to counter innate immune responses within hepatocytes via alternative pathways that do not rely on CypA. Since the cyclophilins operate primarily as prolyl isomerases, it is possible that this function in HEV is served via other cellular cyclophilins such as Cyclophilin D (CypD). Interestingly, HAV has been demonstrated to localise to the mitochondria during hepatocyte cell culture infection, suggesting there could be a link between a lack of CypA dependence and mitochondrial localisation, this is also the site of CypD localisation [35].

In conclusion we suggest that, unlike HCV, HEV is not dependent on CypA to facilitate replication in hepatocytes. We propose that CypB contributes to genotype 1 HEV replication in hepatocytes but is not essential. These observations suggest that the exploitation of CypA by HCV to suppress innate immune responses within hepatocytes, is not required by HEV and that this virus may have other mechanisms to prevent elimination by the innate immune responses at work within hepatocytes.

## Author contributions

FJTB, MRH, and MH designed the study and wrote the manuscript. FJTB conducted the replication and survival assays. SC generated the silenced cell lines used in these experiments. FJTB and MRH analysed the data. MRH and MH provided supervision.

## Acknowledgements

We thank Patrizia Farci (National Institute of Allergy and Infectious Diseases, Bethesda) and Koji Ishii (National Institute of Infectious Diseases, Tokyo) for the genotype 1 and genotype 3 HEV replicons, respectively.

## Materials & correspondence

Correspondence and materials requests should be directed to MRH.

